# Semisynthesis of functional transmembrane proteins in GUVs

**DOI:** 10.1101/2021.09.08.459519

**Authors:** K. A. Podolsky, T. Masubuchi, G. T. Debelouchina, E. Hui, N. K. Devaraj

**Author notes:** Corresponding Author: Neal K. Devaraj – Department of Chemistry and Biochemistry, University of California, San Diego, La Jolla, California 92093, United States.

## Abstract

Cellular transmembrane (TM) proteins are essential sentries of the cell facilitating cell-cell communication, internal signaling, and solute transport. Reconstituting functional TM proteins into model membranes remains a challenge due to the difficulty of expressing hydrophobic TM domains and the required use of detergents. Herein, we use a intein-mediated ligation strategy to semisynthesize bitopic TM proteins in synthetic membranes. We have adapted the trans splicing capabilities of split inteins for a native peptide ligation between a synthetic TM peptide embedded in the membrane of giant unilamellar vesicles (GUVs) and an expressed soluble protein. We demonstrate that the extracellular domain of programmed cell death protein 1 (PD-1), a mammalian transmembrane immune checkpoint receptor, retains its function for binding its ligand PD-L1 at a reconstituted membrane interface after ligation to a synthetic TM peptide in GUV membranes. We envision that the construction of full-length TM proteins using orthogonal split intein-mediated semisynthetic protein ligations will expand applications of membrane protein reconstitution in pharmacology, biochemistry, biophysics, and artificial cell development.

## Main text

Transmembrane (TM) proteins comprise about 23% of the human proteome and conduct essential cellular functions such as solute transport, cell-to-cell communication, signaling, and lipid synthesis.^1,2^ As vital gatekeepers to the cell interior, TM proteins have high therapeutic relevance, and are targeted by about 60% of currently available pharmaceuticals.^3^ Reconstituting TM proteins into model membranes (*e.g*. unilamellar vesicles, supported lipid bilayers, and nanodiscs) enables functional and structural studies of these proteins in near-native environments for biochemical and biophysical analysis.^4–7^ Protein reconstitution into model membranes such as giant unilamellar vesicles (GUVs), relies on expressing and purifying full-length TM proteins in the presence of detergents which are known to unpredictably affect protein function and structure.^7–12^ In practice, expressing and purifying TM proteins is often challenging due to their general hydrophobicity leading to aggregation.^13,14^ Some studies have avoided these pitfalls by chemically lipidating proteins or expressing truncated soluble domains of TM proteins, and later conjugating the domains to synthetic lipids in model membranes to circumvent expressing the hydrophobic TM region.^15–19^ While eliminating the transmembrane domain eases the tedium of hydrophobic protein expression, this strategy is not ideal, as integral membrane proteins are embedded in cellular membranes by TM domains that are highly conserved and play a variety of roles such as lipid environment sensing, controlling lateral diffusion, participating in dimerization, and facilitating transport and signaling across the bilayer.^20–30^ Thus, there is a need improve the synthesis and reconstitution of TM proteins, ideally without the use of detergents.

Here we demonstrate an effective way of tackling this problem through protein semisynthesis, or the construction of a protein from expressed and synthesized segments.^31^ Specifically, we sought a simplified method to synthesize bitopic (single-pass) TM proteins, which represent about 50% of all TM proteins in the human genome.^32,33^ TM domains pose a problem for protein overexpression and purification due to their toxicity in cell membranes at high concentrations and insolubility in aqueous solutions.^34,35^ Instead, chemical synthesis of a protein TM domain avoids this problem and additionally enables unnatural modification of the synthetic peptide (e.g. unnatural amino acids, fluorophores, site-specific isotope labeling, etc). The subsequent formation of a native peptide bond to an expressed protein soluble domain would avoid the many difficulties of protein reconstitution and offer a more ideal functional model compared to protein-lipid conjugations. We adapt a split intein-mediated expressed protein ligation strategy to build a semisynthetic TM protein into GUVs.^36–39^ Split intein expressed protein ligation is advantageous in comparison to other protein semisynthesis strategies as it requires no additional chemical modifications after protein and peptide production.^39–41^ Natural and engineered split inteins facilitate posttranslational protein trans splicing events and have been used for the determination of protein-protein interactions,^38,42^ protein purification,^43^ labeling of proteins with small molecules or unnatural amino acids,^44–48^ segmental isotopic labeling for nuclear magnetic resonance (NMR),^40,49–52^ semisynthetic protein assembly in mammalian cells,^53,54^ and cell membrane recruitment in expressed protein systems^55^ (Figure S1).^37,56^ Our strategy incorporates synthetic TM peptides directly in model cellular membranes, GUVs,^57,58^ which bypasses the necessity of using cellular machinery to build and insert TM domains into a living cell membrane before reconstitution.^47,48^ To establish the utility of the split intein ligation as a method to build TM proteins in an artificial membrane, we ligated an expressed GFP-intein construct and a synthetic intein-TM peptide embedded in GUVs. Furthermore, we demonstrate the versatility of the ligation in GUVs using the expressed extracellular domain of a glycosylated mammalian bitopic TM receptor PD-1, which retains its ligand-binding activity in a commonly used GUV-based functional assay.

We first identified a suitable split intein-mediated ligation system compatible with semisynthetic reactions. The engineered Cfa_GEP_ split intein system, derived from the ultrafast Cfa_WT_, was chosen for its improved extein tolerance which enables versatility in protein semisynthesis.^36^ Cfa_GEP_ is reportedly robust for semisynthesis, contains a small C intein (38 amino acids) ideal for peptide synthesis, and results in minimal amino acid scaring between exteins. Using Cfa_GEP_, we designed a proof-of-concept semisynthetic pair capable of ligating in phospholipid membranes (Figure 1A). Green fluorescent protein (GFP) was chosen as the protein extein, as it is useful for downstream imaging experiments. GFP fused to Cfa^N^ with a C-terminal polyhistidine tag (GFP-Cfa^N^-His_6_) was expressed in *E. coli* and purified by Ni-NTA column (Figure 1B, Figure S2). A well-characterized, single-pass transmembrane (TM) peptide known as a WALP was chosen as a model synthetic TM peptide extein.^59,60^ WALPs classically contain leucine and alanine (LA) repeats flanked by two tryptophans (WW) on each terminus. A Cfa^C^-WALP peptide was produced via solid phase peptide synthesis (SPPS) on a peptide synthesizer (CEM Liberty Blue; Figure 1B). A fluorescent derivative of Cfa^C^-WALP containing a lysine side chain conjugated to carboxyfluorescein (CfaC-WALP-CF) was also synthesized. After purification of the TM peptides on a reverse phase C-18 column, liquid chromatography electrospray ionization time of flight mass spectrometry (LC-ESI-TOFMS) confirmed their purity and mass (Supplementary spectra 1 & 2).

**Figure 1.**
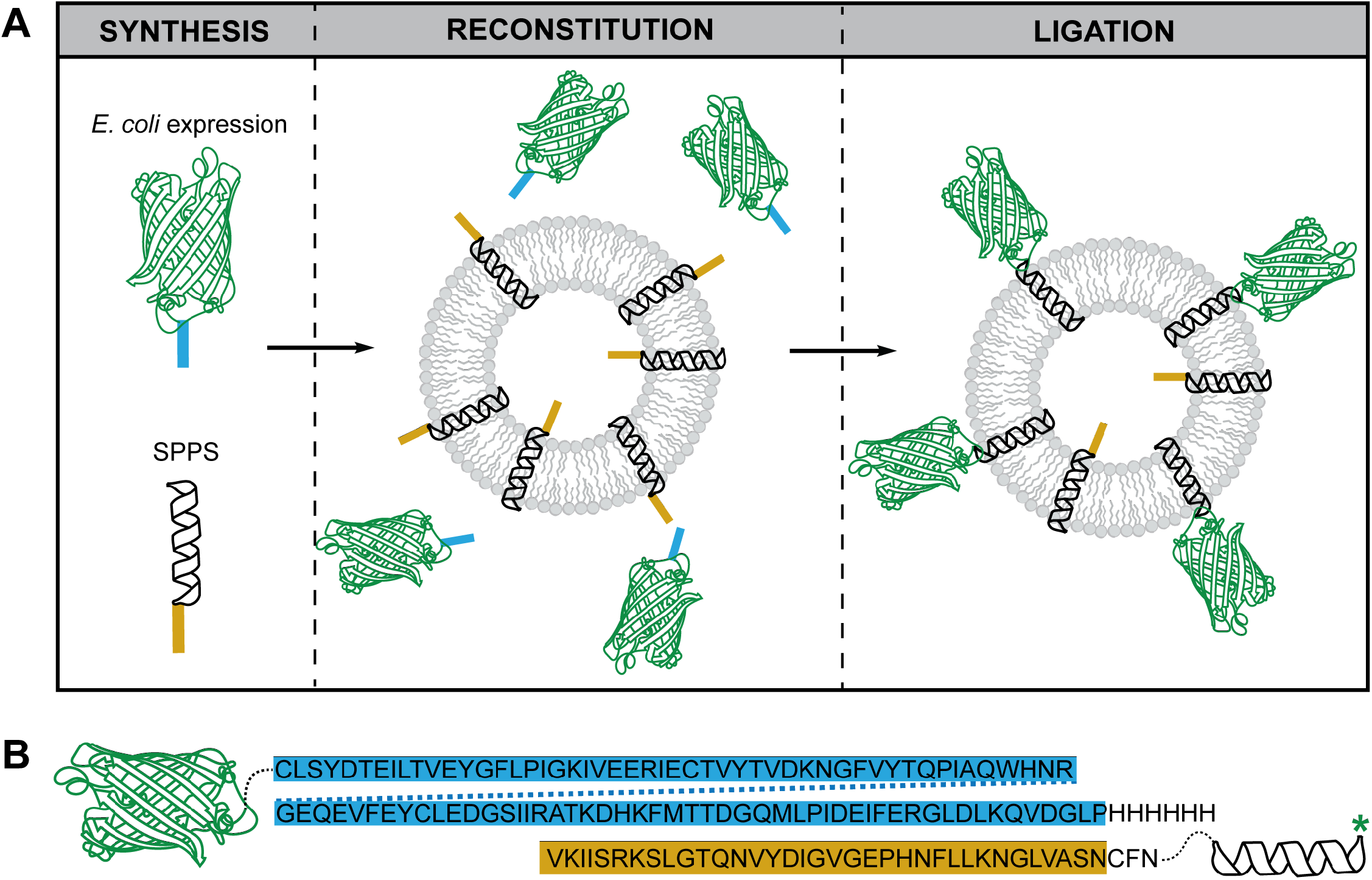
Semisynthetic split intein-mediated ligation. (A) GFP (green), fused to the Cfa^N^ split intein domain (blue) is expressed in *E. coli*, while the WALP (black) fused to the Cfa^C^ split intein domain (yellow) is fabricated via SPPS. Cfa^C^-WALP is reconstituted into GUVs in randomly distributed orientations within the membrane. Upon addition of the soluble GFP-Cfa^N^ construct to peptide-loaded GUVs, the split intein-mediated ligation produces GFP-WALP embedded GUVs. (B) Amino acid sequences of Cfa^N^ (blue) and Cfa^C^ (yellow) inteins and their respective protein and peptide fusions. Dashed lines represent glycine linkers. Green asterisk denotes the position of the CF conjugated to the peptide.

We next verified the reconstitution of TM peptide into phospholipid bilayers. Confocal fluorescence microscopy confirmed the localization of Cfa^C^-WALP-CF to hydrated DOPC vesicles (Figure 2A). Cryo-transmission electron microscopy (cryo-TEM) showed that the vesicles are intact after peptide incorporation and remain intact after the split intein ligation (Figure S3). Circular dichroism (CD) spectra verified that the peptide folds into a secondary alpha helix structure once reconstituted into unilamellar vesicles (Figure 2B, Figure S4).^59–62^ These results show that the synthetic Cfa^C^-WALPs are reconstituting into DOPC bilayers as alpha helical TM peptides.

**Figure 2.**
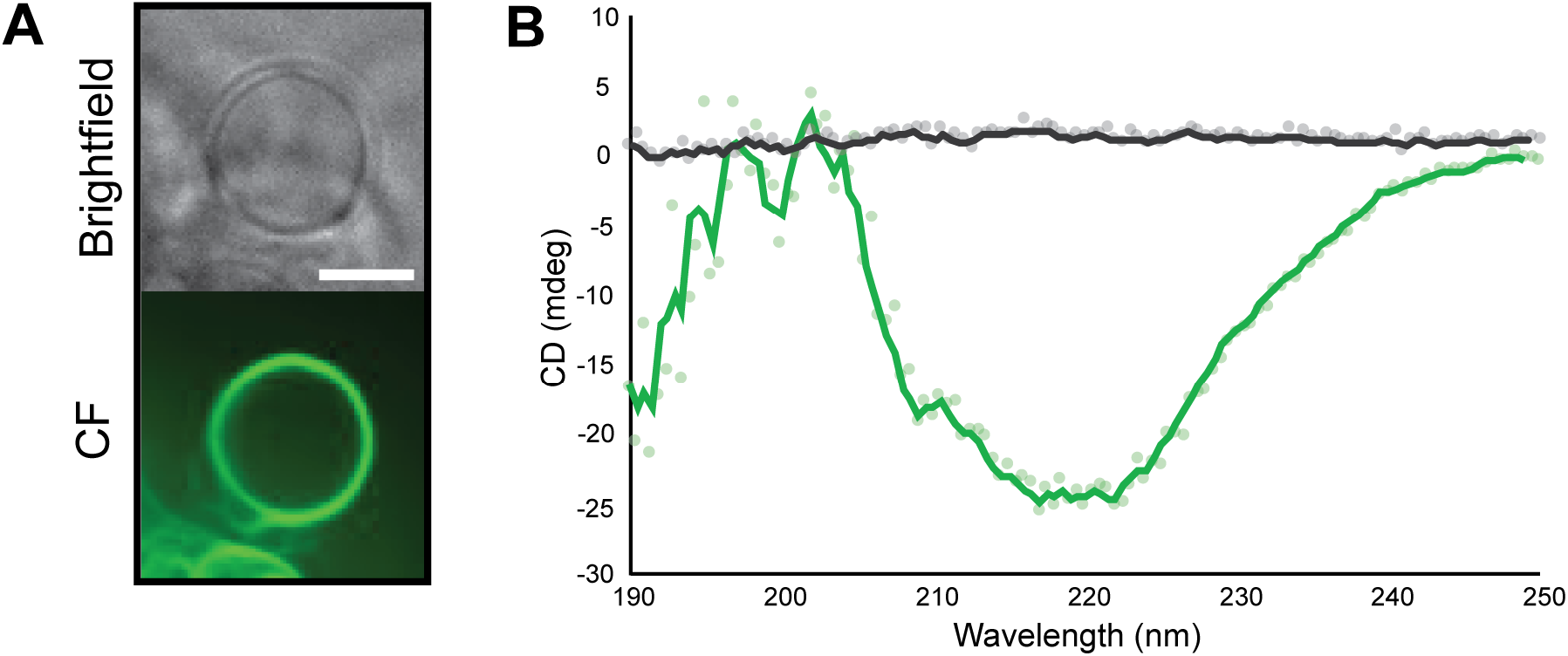
Transmembrane peptide reconstitution. (A) Brightfield and fluorescence (488 nm excitation) microscopy images of a hydrated DOPC vesicle containing Cfa^C^-WALP-CF. Scale bar 10 μm. (B) CD spectra of 20 μM Cfa^C^-WALP before (black) and after (green) reconstitution into SUVs. CF = carboxyfluorescein.

We next proceeded with split intein-mediated GFP-WALP ligation in multilamellar large vesicles (MLVs) formed by hydrating a lipid-peptide film in splicing buffer (150 mM sodium phosphates, 100 mM NaCl, 5 mM EDTA, 1 mM TCEP, pH 7.2). Mimicking previously established conditions for Cfa_GEP_ splicing, we reacted soluble GFP-Cfa^N^-His_6_ to liposome-reconstituted Cfa^C^-WALP (2:1) in splicing buffer.^36^ After 1 and 24 h, LC-ESI-TOFMS analysis confirmed the presence and expected mass of the predicted GFP-WALP product, **F**, which contains an eight amino acid linker (GGCFNGGG) between the GFP and WALP, (Figure 3 A & B, Supplementary spectra 3). SDS-PAGE corroborated these results, where even at 0 min, there is a rapid conversion of all reacting GFP-Cfa^N^-His_6_, **E**, into an intermediate, **H** (Figure 3 C). LC-ESI-TOFMS suggests that **H** is a covalently-linked, branched intermediate of the intein ligation (GFP-Cfa^C^-WALP). The gel band intensities of **H** and **F** show the conversion of **H** to **F** over time (Supplementary Figure 5). By adapting previously established detergent-free electroformation procedures, we successfully reconstituted CfaC-WALP into GUVs in splice buffer (Table S1).^63–69^ Ligation of GFP-Cfa^N^-His_6_ to TM peptide in GUVs was tracked using confocal fluorescence microscopy and observing fluorescence signal at the lipid membrane (Figure 3 D). The GFP construct alone does not nonspecifically bind to vesicles (Figure 3 D). Taken together, the LC-ESI-TOFMS, SDS-PAGE, and confocal fluorescence microscopy data indicate that the semisynthetic split intein-mediated ligation of protein and TM peptide takes place on liposomes.

**Figure 3.**
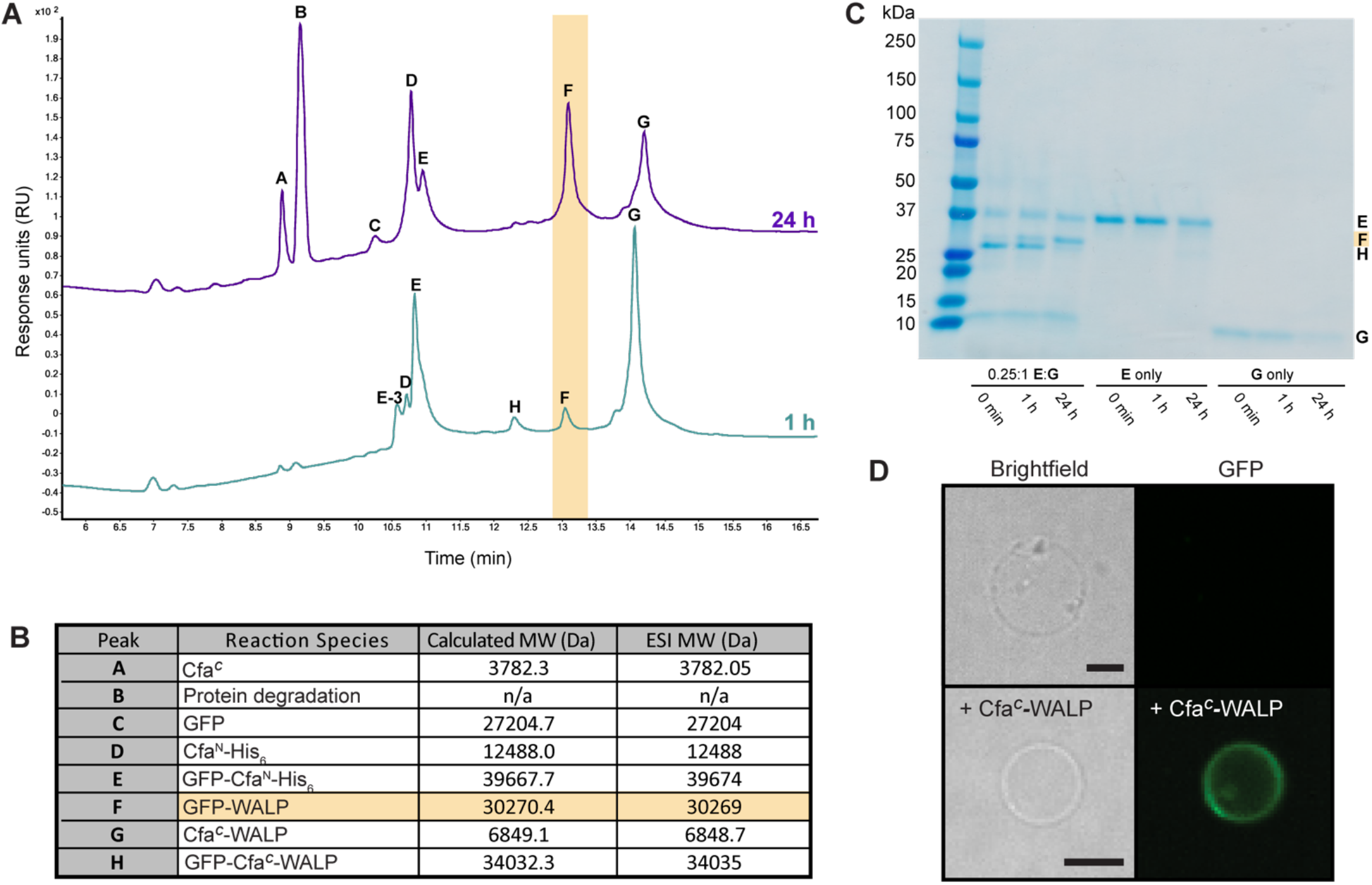
Semisynthetic split intein-mediated ligation to form a transmembrane protein in GUV membranes. (A) Chromatogram of an LC-ESI-TOFMS run of the reaction between GFP-Cfa^N^-His_6_, **E**, and Cfa^C^-WALP, **G**, in vesicles. Each peak corresponds to a reactant, intermediate, or product that is listed in (B) with their corresponding calculated molecular weight (MW) and experimental ESI MW. (C) SDS-PAGE gel of the reaction in (A). Lanes 2-4 are the reaction between **E** and **G**, lanes 5-7 is **E** only, and lanes 8-10 are **G** only. The GFP-WALP product, **F**, is highlighted in yellow boxes throughout the figure. (D) Confocal fluorescence micrographs show that GFP (488 nm excitation) does not bind to GUVs alone (upper row), but does bind to GUVs containing Cfa_C_-WALP after 24 h (lower row). Scale bar, 5 μm.

Membrane proteins are integral to cellular functions and their role in therapeutic intervention is intensively studied. To bypass the challenges of optimizing detergent-mediated reconstitution for functional assays, several studies have turned to tagging extracellular domains of transmembrane proteins and conjugating them to functionalized phospholipids.^15–19^ We chose to focus on PD-1, a transmembrane inhibitory receptor that restricts immune cell function upon binding to its ligand PD-L1 on other cells.^70^ Having been successfully targeted for cancer immunotherapy in a fraction of patients, the PD-1/PD-L1 axis is an urgent focus for translational research.^71^ As with most TM proteins, full length PD-1 is challenging to express and reconstitute into model membranes. Prior effort on membrane reconstitution of PD-1 utilized polyhistidine tags to attach either the intracellular domain or the extracellular domain to a lipid bilayer and thus cannot recapitulate the potential function of the transmembrane domain.^15,29,30,72–75^ Here we sought to reconstitute PD-1 to GUVs via a transmembrane domain using our newly established semisynthesis method. We expressed and purified a fully glycosylated extracellular domain of PD-1 fused to N-terminal SNAP-tag used for fluorescent labeling and C-terminal Cfa^N^ in mammalian HEK293F cells. The recombinant protein was labeled with Janelia Fluor 646 (JF) for downstream fluorescence microscopy experiments and purified using standard procedures outlined in the methods (Figure S6). Although glycosylation of PD-1 resulted in smeared starting material and product bands which are challenging to interpret by SDS-PAGE (Figure S7), the JF-PD-1-Cfa^N^ protein construct was reacted with Cfa^C^-WALP-CF in GUVs and the successful ligation was readily monitored via fluorescence microscopy (Figure 4 A, Figure S7, Figure S8).

**Figure 4.**
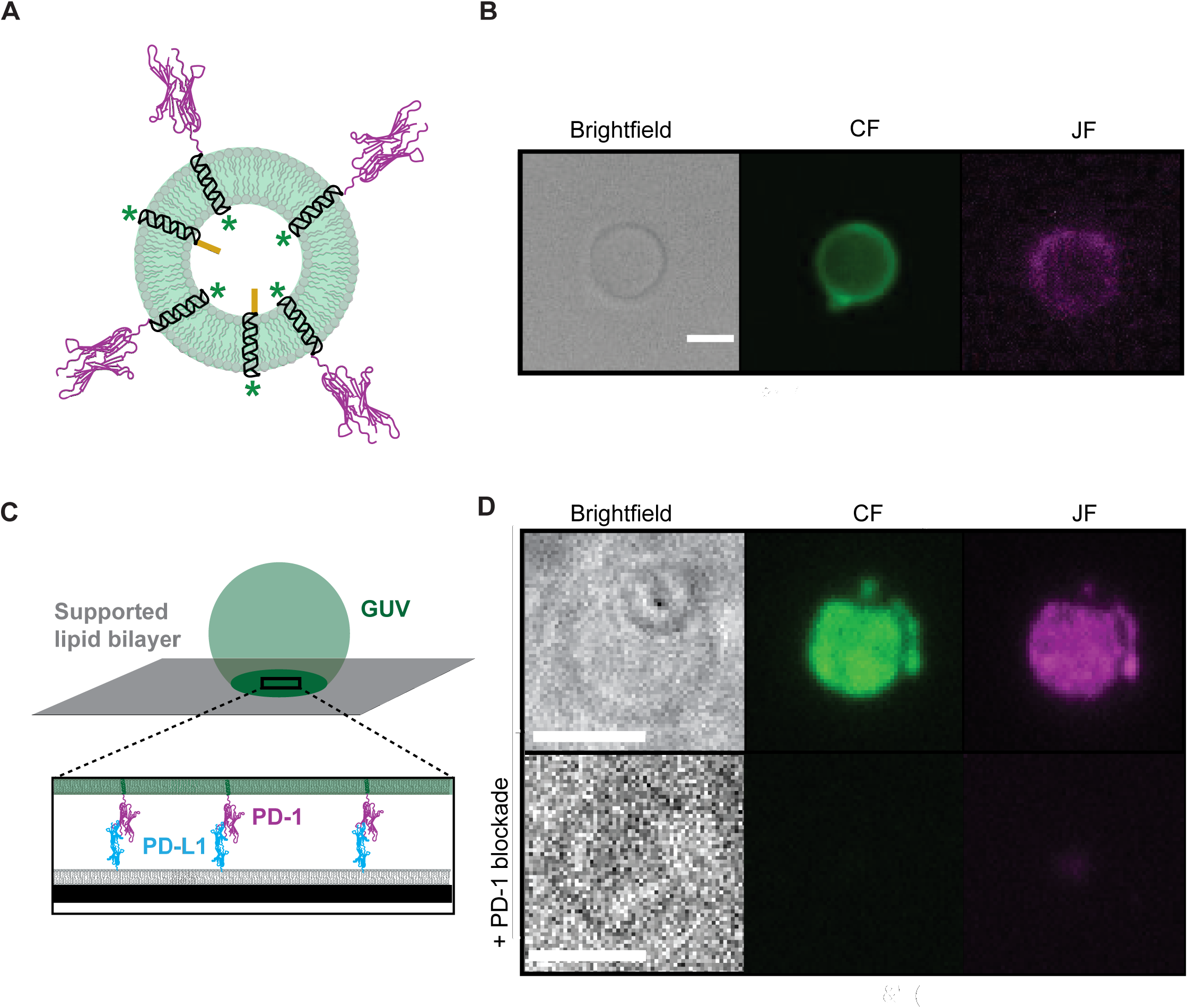
Building a functional semisynthetic transmembrane protein in GUVs. (A) A cartoon representation depicts the CF-labeled (green asterisks) transmembrane peptide (black) fused to the extracellular domain of PD-1-JF (magenta). (B) Brightfield and fluorescent micrographs of the semisynthetic JF-PD-1-WALP-CF transmembrane product in a GUV. CF= 488 nm excitation. JF 646 = 638 nm excitation. (C) Cartoon schematic of the PD-1 activity experiment where a GUV (green) contacts a SLB leading to the enrichment of PD-1 at the GUV-SLB interface due to trans-PD-1:PD-L1 interaction. (D) TIRF brightfield and fluorescence micrographs of the GUV-SLB interface showing enrichment of CF and JF signals at the interface. In the presence of PD-1 blockade (bottom row), there is no enrichment of either signal although a GUV remains at the SLB surface. Because the GUV is unbound to the SLB surface, the GUV is not clearly seen on this focal plane. A mid-GUV cross section better depicts this unbound GUV (Figure S9). CF = carboxyfluorescein. JF = Janelia Fluor 646.

To show that the extracellular domain of PD-1 remains functional after split intein-mediated semisynthesis, we used total internal reflection fluorescence (TIRF) microscopy to visualize trans-PD-1:PD-L1 interaction at a membrane interface between a PD-1 GUV and a PD-L1-attached planar lipid bilayer. To produce PD-1 GUVs, we reacted JF-PD-1-Cfa^N^ with electroformed GUVs containing semisythetic Cfa^C^-WALP-CF in splice buffer. To construct a ligand (PD-L1) presenting membrane, we attached polyhistidine-tagged PD-L1 to a supported lipid bilayer (SLB) doped with 5% NTA-DGS lipids, following standard procedures.^15^ Upon incubating the semisythetic PD-1 GUVs with PD-L1 SLBs, we visualized PD-1 distribution at the GUV-SLB contact area by TIRF fluorescence microscopy (Figure 4 B). As a control, to one well, a PD-1 antibody blockade, pembrolizumab, was added to inhibit the binding of PD-1 to PD-L1. Successful PD-1:PD-L1 trans-interaction would enrich PD-1 to the GUV-SLB interface as previously observed.^72^ Indeed, we found that, in the absence of pembrolizumab, the PD-1 ectodomain and TM peptide fluorescent signals are co-enriched at the SLB-GUV interface, evidence for successful ligation and for the ability of the semisythetic PD-1 to engage PD-L1 in trans. In the presence of PD-1 blockade, the sunken GUV was unable to bind but remained in close proximity to the SLB, indicated by the bright field image of the bottom of the GUV and the minor PD-1 fluorescent signal (Figure S9). These results indicate that semisynthetic PD-1 retains its function after ligation to TM peptides within GUV membranes.

In summary, we have established that split intein-mediated ligations can be used to construct semisynthetic TM proteins in artificial phospholipid membranes, including GUVs. As demonstrated by the trans-PD-1:PD-L1 PD-L1 interaction at the GUV-SLB interface, the semisynthetic TM proteins are functional after ligation. Given the published versatility of the Cfa_GEP_ and other split intein ligations, there should be flexibility to diversify the semisynthetic protein products.^36^ Future work in our lab will demonstrate the use of synthetic native TM sequences for the detergent-free intein-mediated ligation in GUVs. While this initial study focused on ligating only the extracellular domain of a bitopic membrane protein, we plan to leverage the orthogonality of split inteins to semisynthesize full-length, single-pass TM proteins by incorporating an additional intein-mediated ligation site on the C-terminus of the TM peptide that will ligate to an expressed intracellular domain.^76^ We envision that this method will improve biochemical, biophysical, and pharmacological research on TM proteins as it will expedite the production of reconstituted proteins anchored by synthetic or native TM domains in GUVs. We plan to apply our method of split intein-mediated TM protein semisynthesis to study TM protein localization in phase separated GUVs, the reconstruction of signaling networks within artificial cells, and the effects of therapeutic intervention on protein structure and function.

## Author Contributions

The manuscript was written through contributions of all authors.

## Notes

The authors declare no competing financial interest.

## Acknowledgement

The authors thank the UCSD Mass Spectrometry Facility and UCSD Cryo-EM facility for data collection and training. We additionally thank Prof. Akif Tezcan (University of Califor-nia, San Diego) CD training and use. Thank you to J. R. Winnikoff for critical reading of the manuscript.

