## Supplementary Information for "Semisynthesis of functional transmembrane proteins in GUVs"

### Table of Contents

**1. Materials, General Methods, and Instrument Details**
**2. Experimental Procedures**
**3. Supplementary Sequences and Structures**
**4. Supplementary Tables and Figures**
**5. LCMS spectra**
**6. References**

### 1. Materials, General Methods, and Instrument Details

Commercially available 1,2-dioleoyl-sn-glycero-3-phosphocholine (DOPC), 1-palmitoyl-2-oleoyl-sn-glycero-3-phosphocholine (POPC) (#850457), 1,2-dioleoyl-sn-glycero-3-[(N-(5-amino-1-carboxypentyl) iminodiacetic acid) succinyl] (nickel salt, DGS-NTA-Ni) (#790404), and 1,2-dioleoyl-sn-glycero-3-phosphoethanolamine-N-[methoxy(polyethylene glycol)-5000] (PEG5000-PE) (#880230) was obtained from Avanti® Polar Lipids. Fmoc-Ala-OH, Fmoc-Arg(Pbf)-OH, Fmoc-Asn(trt)-OH, Fmoc-Asp(OtBu)-OH, Fmoc-Cys(trt)-OH, Fmoc-Gln(trt)-OH, Fmoc-Glu(OtBu)-OH, Fmoc-Gly-OH, Fmoc-His(trt)-OH, Fmoc-Ile-OH, Fmoc-Leu-OH, Fmoc-Lys(Boc)-OH, Fmoc-Phe-OH, Fmoc-Pro-OH, Fmoc-Ser(tBu)-OH, Fmoc-Thr(tBu)-OH, Fmoc-Trp(Boc)-OH, Fmoc-Tyr(tBu)-OH, and Fmoc-Val-OH were purchased from ChemImpex. Fmoc-Lys(5/6-FAM)-OH was purchased from AnaSpec. N,N-dimethylformamide (DMF), acetonitrile (ACN), N,N-diisopropylethylamine (DIEA), trifluoroacetic acid (TFA), triisopropylsilane (TIS), 2-2'-(ethylenedioxy)diethanethiol (DODT), N,N'-diisopropylcarbodiimide (DIC), tris(2-carboxyethyl)phosphine hydrochloride (TCEP), 4-methylpiperidine, chloroform, anhydrous dichloromethane (DCM), anhydrous diethylether, and anhydrous methanol (MeOH) were obtained from Sigma-Aldrich. Oxyma was purchased from CEM. JaneliaFluor646-SNAP ligand was provided from Janelia Research Campus (HHMI) (#Janelia 2014-013). Human PD-L1-His recombinant protein was purchased from Sino Biological (#10084-H08H). Pembrolizumab was purchased from Selleck Chemicals (#A2005). HEK293F cells were obtained from Dr. Andrew Ward (Scripps Research). All reagents obtained from commercial suppliers were used without further purification unless otherwise noted.

Spinning-disk confocal microscopy images were acquired on a Yokagawa spinning disk system (Yokagawa, Japan) built around an Axio Observer Z1 motorized inverted microscope (Carl Zeiss Microscopy GmbH, Germany) with a 63x, 1.40 NA oil immersion objective or 20x 0.8 NA objective to an ORCA-Flash 4.0 V2 Digital CMOS camera (Hamamatsu, Japan) using ZEN Blue imaging software (Carl Zeiss Microscopy GmbH, Germany). The fluorophores were excited with diode lasers (405 nm-20 mW, 488 nm-30 mW, 561 nm-20 mW, and 638 nm-75 mW). A condenser/objective with a phase stop of Ph3 was used to obtain the phase-contrast images with a 20x objective on an Olympus BX51 upright fluorescent microscope. The fluorophores were excited with 20 mW DPSS lasers (GFP, JF 646). For cryo-TEM analysis, vesicles were imaged on a Titan Krios G3 transmission electron microscope (ThermoFisher) operated at 300 kV with an energy filter (Gatan), and volta phase plates. Images were recorded on a K2 Summit direct electron detector (Gatan). Images were taken at 33,000 x nominal magnification with a 4.33 angstrom pixel size and calibrated 11,575 x magnification. HPLC purification was carried out on Zorbax SB-C18 semipreparative column with Phase A/Phase B gradients [*Phase A*: H2O with 0.1% v/v TFA; *Phase B*: ACN with 0.1% v/v TFA] and monitored via diode array detector at 210 nm. Electrospray Ionization-Mass spectra (ESI-MS) were obtained on an Agilent 6230 Accurate-Mass Time of Flight (TOF) mass spectrometer from the UCSD Mass Spec Facility. Circular dichroism (CD) experiments were done on an AVIV CD Spectrometer Model 215. Scans were taken of each sample from 190-250 nm with a 1.0 nm Bandwidth in triplicate at room temperature.

### 2. Experimental Procedures

Cloning GFP-$Cfa^N$-$His_6$

pTXB1, containing the $Cfa^N$-$His_6$ split intein gene, was kindly provided to us by Professor Tom Muir's lab at Princeton University. A pET-11a cloning vector containing the sequence of the sfGFP construct was provided by Dr. Henrike Niederholdtmeyer in Dr. Neal Devaraj's lab. PCR and isothermal assembly (NEBuilder, New England Biolabs) was used per the vendor's instructions to insert the split intein gene into pET-11a, yielding a GFP-$Cfa^N$-$His_6$ fusion construct which was transformed into DH5α *E. coli* competent cells. A double glycine linker was placed between GFP-$Cfa^N$ for improved splicing efficiency. After overnight culture at 37 ˚C and 200 rpm, plasmid minipreps were performed (Qiagen) and construct sequence was verified by Sanger Sequencing (Eton Biosciences).

Expression and purification of GFP-$Cfa^N$-$His_6$

Plasmids confirmed to have the correct fusion construct sequence were transformed into BL21 (DE3) *E. coli* competent cells (New England Biolabs) per vendor instructions. These cells were then grown overnight at 37 ˚C in Luria-Bertani (LB) broth containing 0.1 mg/mL carbenicillin, a more stable substitute antibiotic of ampicillin. 1

mL of the overnight culture was used to inoculate 100 mL of autoclaved LB medium containing 0.1 mg/mL carbenicillin. The culture was grown at 37 ˚C with shaking at 200 rpm until the $OD_{600}$ of the culture reached 0.6. Overexpression of GFP-$Cfa^N$-$His_6$ was induced with 0.5 mM isopropyl 1-thio-D-galactopyranoside (IPTG). The cells were then grown for 4 h at 37 ˚C with shaking at 200 rpm and subsequently harvested via centrifugation at 4000 rcf for 20 min at 4 ˚C. The visibly bright green (indicating the presence of full-length GFP) pellet was stored at -80 ˚C until further use. Buffers were prepared as followed: buffer A (50 mM phosphates, 300 mM NaCl, 5 mM imidazole, pH 7.5), wash buffer I (50 mM phosphates, 300 mM NaCl, 20 mM imidazole, pH 7.5), wash buffer II (50 mM phosphates, 200 mM NaCl, 50 mM imidazole, pH 7.5), elution buffer (50 mM phosphates, 300 mM NaCl, 250 mM imidazole, pH 7.5). Cell pellets were thawed and resuspended in lysis buffer (5 mL buffer A, 1 mM PMSF in ethanol) on ice. The resuspended cells were lysed on ice by ultrasonication (35% amplitude for 3 minutes 50% duty cycle with 40 second period at power level 6). The visibly bright green supernatant was incubated in a gravity column containing $Ni^{2+}$-nitrilotriacitate (NTA) resin pre-equilibrated with 10 mM imidazole on a shaker for 1 h at 4 ˚C. A small sample of flow through was kept for SDS-PAGE while the rest was discarded. The resin was washed four times on ice with 2 column volumes (CV; 600 µL) of wash buffer I and two times with 1 CV of wash buffer II by centrifuging the column for 2 seconds at 600 rcf into prepared tubes for collection of the supernatant. The column was washed six times on ice with 200 µL of elution buffer by gravity, each time collecting the visibly bright green eluent fraction in separate Eppendorf tubes. The fractions were analyzed by SDS-PAGE to check for considerable impurities. Fractions were pooled and aliquoted into high and low concentration samples to final concentrations of 19 µM and 373.5 µM. LC-ESI-TOFMS corroborated the purity and verified the correct mass of the protein construct.

SDS-PAGE

All SDS-PAGE experiments were run for 35 minutes at 200 V on 15 well 4-20% MiniPROTEAN TGX Precast Protein Gels (Bio-Rad). Sample was added to loading dye (1:1) at specified time points, then placed on a 95 ˚C heat block for 5 minutes, placed on ice, quickly spun down via tabletop centrifuge, and loaded onto the gel. Gels were stained with Instant Blue Coomassie Stain (Abcam) for 1-24 h and destained with water. Gels were imaged on a tabletop scanner.

Solid phase peptide synthesis (SPPS) of TM peptide

$Cfa^C$-WALP and $Cfa^C$-WALP-CF were synthesized in house via SPPS on a Liberty Blue peptide synthesizer (CEM) at a 0.1 mmol synthesis scale. A triple glycine linker was placed between $Cfa^C$-WALP in both constructs for improved splicing efficiency. Prior to synthesis, amino acids ( 0.2 M in DMF), coupling agent (20% v/v solution of 4-methylpiperidine in DMF), wash solvent (DMF), activator (0.5 M DIC in DMF), and base activator (0.5 M Oxyma and 0.1 eq DIEA in DMF), and pre-loaded resin were prepared. To pre-load the resin, after swelling in anhydrous DCM for 10 minutes, 0.2 g Trityl-OH resin (ChemMatrix) was activated in 3 M acetyl chloride in DCM for 3 min at room temperature with shaking. The resin (0.5 mmol/g loading capacity) was then washed with anhydrous DCM (3 x 3 mL) and an amino acid solution containing 4 eq Fmoc-Ala-OH and 4 eq DIPEA in 2 mL DCM was added. The resin was shaken at room temperature overnight. It was drained and a capping solution of DCM/MeOH/DIEA (17:2:1) was added for 5 min with shaking at room temperature. The resin was then washed (3 x 2 CV of DCM, 2 x 2 CV of DMF, and 3 x 2 CV of DCM) and put on a desiccator to dry until placed in a 30 mL Liberty Blue reaction vessel for synthesis. Resin loading for both peptides were calculated to be ~0.4 mmol/g resin using standard UV absorption method upon Fmoc cleaving of small aliquots of loaded resin.[1] Subsequent protected amino acid couplings were done on the Liberty Blue peptide synthesizer using standard microwave-assisted deprotection and coupling settings. The 20 N-terminal amino acids were double-coupled to ensure coupling to the long-sequence, hydrophobic peptide. After synthesis, the peptide-conjugated resin was removed from the coupling vessel, washed with 3 x 2 CV of DCM, covered with foil, and desiccated overnight. To prepare the fluorescent $Cfa^C$-WALP-CF peptide (rather than the nonfluorescent $Cfa^C$-WALP), the C-terminal standard protected lysine was replaced with Fmoc-Lys(5/6-FAM)-OH (AnaSpec) during synthesis.

Deprotection and purification of TM peptide

Peptide-conjugated resin was shaken for 2 h at room temperature in a 6 mL TFA/TIS/$H_2O$/DODT deprotection solution (37:1:1:1). The filtrate was collected in 15 mL Falcon tubes. Ice cold diethylether was added to each Falcon tube to precipitate the crude peptide product. The tubes were centrifuged at 7500 rcf for 5 min and the supernatant was discarded. The pellet was resuspended in ice cold anhydrous diethylether, centrifuged, and the supernatant was removed two additional times. The pellet was desiccated for 30 min and then dissolved in 0.5 mL $H_2O$/methanol (1:1) and transferred to a weighed glass vial. To the dissolved crude peptide, 1 mL of $H_2O$ was added and the peptide solution was frozen at -80 ˚C for lyophilization overnight. The lyophilized peptide powder was resuspended in $H_2O$/MeOH (1:1) for HPLC purification (Zorbax SB-C18 semipreparative column, 5% v/v $H_2O$ + 0.1% v/v TFA in ACN + 0.1% v/v TFA; 10-11 min). The purified fraction was concentrated, lyophilized, and obtained as a white or yellow powder for $Cfa^C$-WALP and $Cfa^C$-WALP-CF, respectively. For experiments, 200 µM purified TM peptide stock solutions were freshly prepared in chloroform, vials sealed with parafilm, and stored at -20 ˚C for up to two weeks.

Reconstitution of TM peptide in multilamellar vesicles (MLVs)

To reconstitute TM peptides into MLVs, a hydration method for vesicle formation was used. DOPC (25 µL, 10 mM) and TM peptide (25 µL, 200 µM) were mixed (50:1 lipid/peptide) and dried into a lipid and TM peptide film by $N_2$ gas stream in a glass scintillation vial. The vial was desiccated for 30 min. Water or splice buffer (250 µL) was added to the vial which was then rotated at room temperature for 1 hour and vortexed. Confocal microscopy verified the formation of vesicles and the localization of TM fluorescent peptide to the MLVs.

Reconstitution of TM peptide in GUVs

Large, unilamellar artificial bilayers recapitulate the structure and curvature of natural cell membranes to model cellular processes, so we sought to bring this ligation system into GUVs. We turned to electroformation as a common method of GUV formation that does not use protein structure- and function-altering detergents. Unfortunately, common electroformation methods using indium tin oxide-coated glass slides are incompatible with high salt buffer such as the splicing buffer essential to carry out the intein-mediated ligation reaction in GUVs.[2] It was necessary to turn to alternative simultaneous electroformation and reconstitution methods compatible with the splice buffer. There has been success forming GUVs with reconstituted membrane proteins in high salt buffer using platinum (Pt) wires and sequential changes in voltage sine wave parameters (Table S1).[3–9] We adapted these previously published methods for GUV formation in high salt buffers to reconstitute TM peptides into GUVs. First, hydrated vesicle samples of a 50:1 lipid/peptide ratio were prepared in water as described above. These were sonicated in a bath sonicator for 1 h to form SUVs. In a Pt wire electroformation device compatible with confocal microscopy, seven drops of SUVs were placed on each Pt wire. The device was placed at 40 ˚C until the droplets were dried (~5 min). Seven drops of SUVs were placed on each Pt wire and dried again at 40 ˚C for ~5 min. The device was placed in a 40 ˚C chamber either on or off of the confocal microscope stage for monitoring the electroformation. Splice buffer (800 µL) was added right before a 60 MHz DDS signal generator (Koolerton) applied specific voltage and frequency to the electroformation device over a 3.5 h time period (Supplementary Table 1). After electroformation, GUVs were gently lifted off by gently tapping the device and pipetting splice buffer from inside the device over the wires 10 times. Before pipetting, 1 cm of the pipette tip was snipped off with scissors to prevent GUV collapse due to shearing forces within the smaller opening of the tip. The detached GUVs were visualized by spinning-disk confocal or compound microscopy.

Cryo-TEM of vesicles with reconstituted TM peptide

Cryogenic transmission electron microscopy (cryo-TEM) was used to verify that there was no disruption of the lipid membranes by peptide incorporation and no visible accumulation of peptide at vesicle surfaces indicating its reconstitution into DOPC membranes (Figure S3). MLVs with and without TM peptide reconstitution were prepared as described above. MLVs were sonicated in a bath sonicator for 1 h. Immediately before grid vitrification, the sample was pipetted onto plasma-cleaned 200-mesh Quantifoil R 2/2 copper grids (Quantifoil). Using a Vitrobot EM grid plunger (FEI), excess buffer was blotted at room temperature and 95% humidity and the grids were plunge-frozen in liquid ethane maintained at −180 °C. The grids were stored in liquid nitrogen until use. After clipping under cryogenic conditions and placing them in the autoloader cassette, the grids were loaded onto a Titan Krios for data collection using EPU (ThermoFisher) and DigitalMicrograph (Gatan) softwares.

Circular Dichroism

We adapted previous methods for analyzing the folding of WALPS reconstituted into lipid membranes via CD.[10,11] Briefly, TM peptide reconstituted in SUVs were prepared reconstituting TM peptide in MLVs in water as described above (30:1 lipid/peptide ratio) and ultrasonicating the MLV sample for 3 minutes on ice (40% amplitude, power level 6). The SUV samples were spun down at 15000 rpm at 24 ˚C for 5 minutes and the supernatant was used for CD measurement. TM peptide samples without DOPC present were prepared by making a peptide film and hydrating the sample in water (final concentration 20 μM). Samples and a water blank were run in triplicate on the CD spectrometer. For data analysis, all samples and blanks were averaged. The data was baseline corrected by subtracting the water blank average from the peptide-containing sample averages. The baseline corrected averages were plotted and a moving average trendline with a period of three was added to the data for clarity.

Expression and purification of JF-PD-1-Cfa$^N$

The ectodomain of human PD-1 (aa 24-170) with an N-terminal signal peptide of HIV envelop glycoprotein gp120 followed by a SNAP-tag, and with a C-terminal Cfa$^N$ followed by a TwinStrep-tag (PD-1-Cfa$^N$) was cloned into a pPPI4 plasmid and expressed in HEK293F cells as described previously.[12,13] The secreted proteins were purified through StrepTrap HP column (GE Healthcare, 28907547) and labeled with JaneliaFluor646-conjugated SNAP ligand (JF, Janelia research). The labeled monomeric proteins were further purified using a Superdex 200 increase 10/300 GL column (GE Healthcare, 28990944) in HEPES buffered saline (50 mM HEPES, pH 7.5, 150 mM NaCl, 10% glycerol). The purified protein was quantified by SDS-PAGE and Coomassie blue staining using bovine serum albumin (BSA, Thermo Scientific, 23209) as a standard, and stored at -80 °C until use. Human PD-L1 protein with a C-terminal His-tag was purchased from Sino Biological (10084-H08H).

Supported Lipid Bilayer (SLB) Preparation

Supported lipid bilayers (SLBs) were prepared as described previously (Zhao et al., 2019) with slight modification. A glass-bottomed 96-well plate (Cellvis, P96-1.5H-N) was cleaned with 2.5% Hellmanex (Sigma, Z805939) overnight followed by extensive wash with ddH2O. The washed plate was dried with N2 gas, sealed and stored at room temperature until use. Right before use, wells were etched with 6 M NaOH at 50 °C for 1.5 hours and washed with ddH2O and PBS. SUVs (97.9% POPC, 2% DGS-NTA-Ni, and 0.02% PEG5000-PE) were prepared as previously described and added to the cleaned wells with 100 μL PBS.[14] The wells were incubated at 50 °C for 2 hours and at room temperature for 30 minutes to form SLBs. The excess SUVs were removed by washing with PBS and the SLBs were functionalized with 3 nM PD-L1-His protein at room temperature for 1 hour. The unbound PD-L1 was removed by washing with PBS and the wells were equilibrated with GUV imaging buffer (100 mM Sodium phosphate, 150 mM NaCl, 1 mM EDTA, 100 mM glucose, pH 7.2).

TIRF Microscopy of GUV-SLB Contact

The JF-PD-1-WALP-CF reconstituted GUVs were mixed with or without 40 μg/mL Pembrolizumab and incubated at RT for 10 minutes, and added to the SLB-containing wells with 100 μL GUV imaging buffer. The wells were incubated at room temperature for 10 minutes to let the GUVs settle on the SLB. The fluorescence of GREEN fluorophore and PD-1*JF646 were visualized using Nikon Eclipse Ti TIRF microscope equipped with a 100x Apo TIRF 1.49 NA objective, controlled by the Micro-Manager software.[15] Images were processed using Fiji.9/8/21 5:38:00 PM

Semisynthesis of GFP-WALP and JF-PD-1-WALP-CF in MLVs

Because lipid film hydration can produce multilamellar vesicles at high lipid and peptide concentrations that are necessary for MS and SDS-PAGE analysis, MLVs were used for those experiments. Expressed protein (GFP or PD-1) was diluted in splice buffer to a final concentration of 5 μM and left on ice for 5 minutes. To start the reaction, TM peptide reconstituted into MLVs in splice buffer (40 μL) was added to the expressed protein solution. The reaction was put on a rotator at 37 ˚C for up to 24 h and samples were sometimes taken from the solution to monitor the reaction progress via SDS-PAGE (8 μL) or LC-ESI-MS (100 μL). Samples could then be visualized by spinning-disk confocal or compound microscopy.

Semisynthesis of GFP-WALP and JF-PD-1-WALP-CF in GUVs

Expressed protein (GFP-Cfa$^{N}$ or JF-PD-1-Cfa$^{N}$) was added to Cfa$^{C}$-WALP-containing GUVs electroformed in splice buffer (described above) to a final concentration of 1 µM. The reaction was put on a rotator at 37 ˚C for 24 h. Samples were then visualized by spinning-disk confocal or compound microscopy.

#### 3. Supplementary Sequences and Structures

Single-letter amino acid sequences of synthetic peptide and expressed protein constructs. Cfa$^{C}$ (blue), Cfa$^{N}$ (yellow), linker (red), CF (green), and SNAP-tag (purple) are highlighted.

Cfa$^{C}$-WALP

VKIISRKSLGTQNVYDIGVGEPHNFLLKNGLVASNCFNGGGWWLALALALALALALALALWWKA

Cfa$^{C}$-WALP-CF

VKIISRKSLGTQNVYDIGVGEPHNFLLKNGLVASNCFNGGGWWLALALALALALALALALWWKA

GFP-Cfa$^{N}$-His$_{6}$

MKSSRKGEELFTGVVPILVELDGDVNGHKFSVRGEGEGDATNGKLTLKFICTTGKLPVPWPTLVTTLTYGVQCFARYPDHM
KQHDFFKSAMPEGYVQERTISFKDDGTYKTRAEVKFEGDTLVNRIELKGIDFKEDGNILGHKLEYNFNSHNVYITADKQKN
GIKANFKIRHNVEDGSVQLADHYQQNTPIGDGPVLLPDNHYLSTQSVLSKDPNEKRDHMVLLEFVTAAGITHGMDELYKGG
CLSYDTEILTVEYGFLPIGKIVEERIECTVYTVDKNGFVYTQPIAQWHNRGEQEVFEYCLEDGSIIRATKDHKFMTTDGQM
LPIDEIFERGLDLKQVDGLPHHHHHH

GFP-WALP

MKSSRKGEELFTGVVPILVELDGDVNGHKFSVRGEGEGDATNGKLTLKFICTTGKLPVPWPTLVTTLTYGVQCFARYPDHM
KQHDFFKSAMPEGYVQERTISFKDDGTYKTRAEVKFEGDTLVNRIELKGIDFKEDGNILGHKLEYNFNSHNVYITADKQKN
GIKANFKIRHNVEDGSVQLADHYQQNTPIGDGPVLLPDNHYLSTQSVLSKDPNEKRDHMVLLEFVTAAGITHGMDELYKGG
CFNGGGWWLALALALALALALALALWWKA

Signal peptide-SNAP-PD-1-Cfa$^{N}$-TwinStrep

MDAMKRGLCCVLLLCGAVFVSPSQEIHARFREFMDKDCEMKRTTLDSPLGKLELSGCEQGLHEIKLLGKGTSAADAVEVPA
PAAVLGGPEPLMQATAWLNAYFHQPEAIEEFPVPALHHPVFQQESFTRQVLWKLLKVVKFGEVISYQQLAALAGNPAATAA
VKTALSGNPVPILIPCHRVVSSSGAVGGYEGGLAVKEWLLAHEGHRLGKPGLGGSGSGSFLDSPDRPWNPPTFSPALLVVT
EGDNATFTCSFSNTSESFVLNWYRMSPSNQTDKLAAFPEDRSQPGQDCRFRVTQLPNGRDFHMSVVRARRNDSGTYLCGAI
SLAPKAQIKESLRAELRVTERRAEVPTAHPSPSPRPAGQFQTLVGGCLSYDTEILTVEYGFLPIGKIVEERIECTVYTVDK
NGFVYTQPIAQWHNRGEQEVFEYCLEDGSIIRATKDHKFMTTDGQMLPIDEIFERGLDLKQVDGLPEFWSHPQFEKGGGSG
GGSGGSAWSHPQFEK

Signal peptide-SNAP-PD-1-WALP-CF

MDAMKRGLCCVLLLCGAVFVSPSQEIHARFREFMDKDCEMKRTTLDSPLGKLELSGCEQGLHEIKLLGKGTSAADAVEVPA
PAAVLGGPEPLMQATAWLNAYFHQPEAIEEFPVPALHHPVFQQESFTRQVLWKLLKVVKFGEVISYQQLAALAGNPAATAA
VKTALSGNPVPILIPCHRVVSSSGAVGGYEGGLAVKEWLLAHEGHRLGKPGLGGSGSGSFLDSPDRPWNPPTFSPALLVVT
EGDNATFTCSFSNTSESFVLNWYRMSPSNQTDKLAAFPEDRSQPGQDCRFRVTQLPNGRDFHMSVVRARRNDSGTYLCGAI
SLAPKAQIKESLRAELRVTERRAEVPTAHPSPSPRPAGQFQTLVGGCFNGGGWWLALALALALALALALALWWKA

### 4. Supplementary Tables and Figures

**Table S1.** Electroformation sine wave parameters using Pt wires in splice buffer.

| Step | Voltage (V) | Frequency (Hz) | Time |
|---|---|---|---|
| 1 | 0.106 | 500 | 5 |
| 2 | 0.940 | 500 | 20 |
| 3 | 2.61 | 500 | 185 |

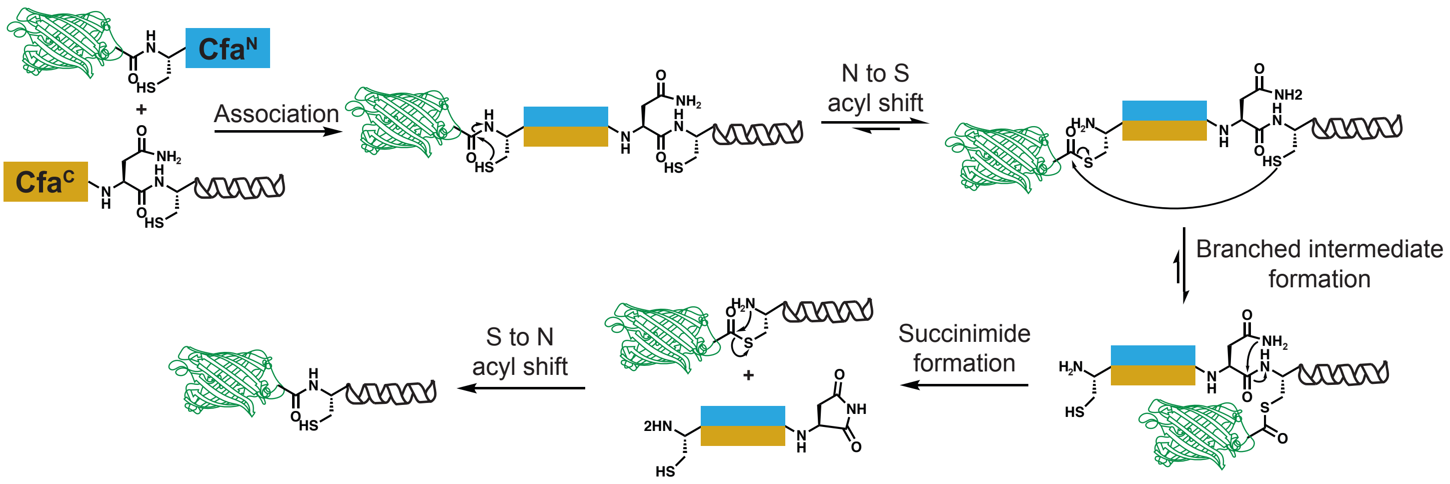


**Figure S1.** Reaction scheme of the general mechanism of the split intein-mediated protein ligation (protein trans-splicing events). The Cfa domains are shown in blue and yellow. GFP is a green cartoon and the WALP is depicted as a black cartoon alpha helix. The product is a native peptide bond between GFP and WALP.

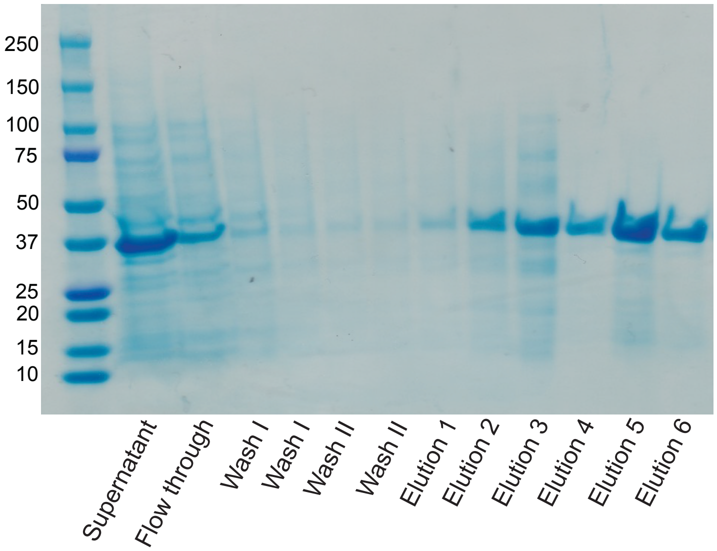


**Figure S2.** Scan of a SDS-PAGE gel of the $Ni^{2+}$-NTA affinity binding purification of GFP-$Cfa^{N}$-$His_6$ (39.7 kDa).

A

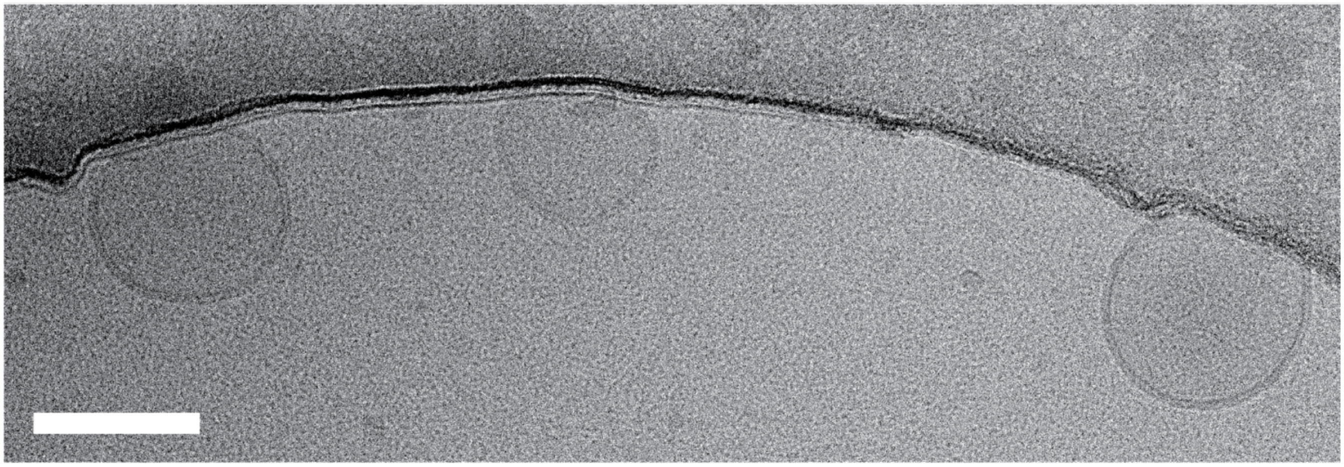

B

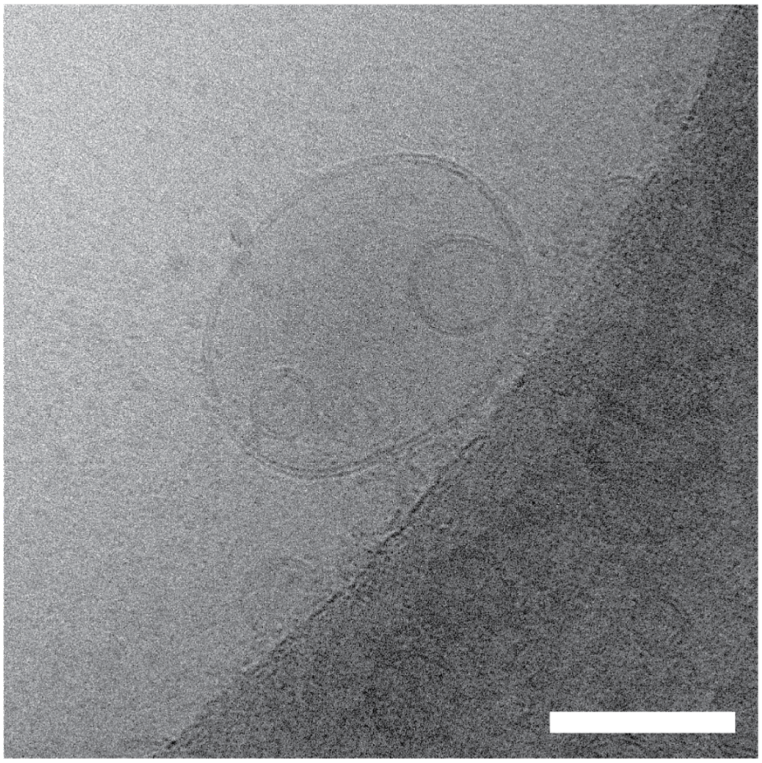

**Figure S3.** Cryo-TEM images of WALP-reconstituted DOPC liposomes (A) and GFP-WALP-reconstituted DOPC liposomes (B). Scale bar, 50 nm.

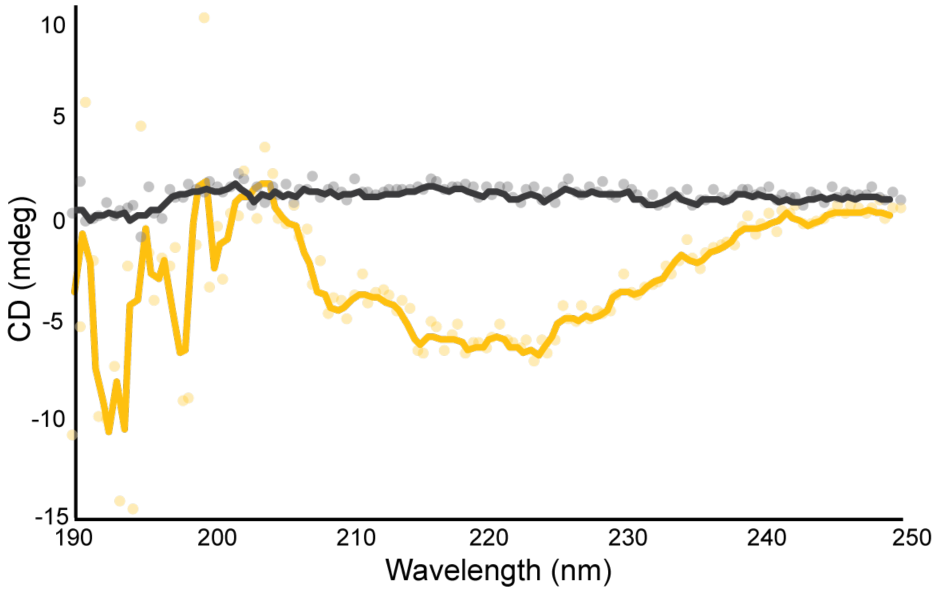


**Figure S4.** CD spectra of DOPC SUVs reconstituted with $Cfa^C$-WALP-CF (yellow) and $Cfa^C$-WALP-CF peptide alone (black).

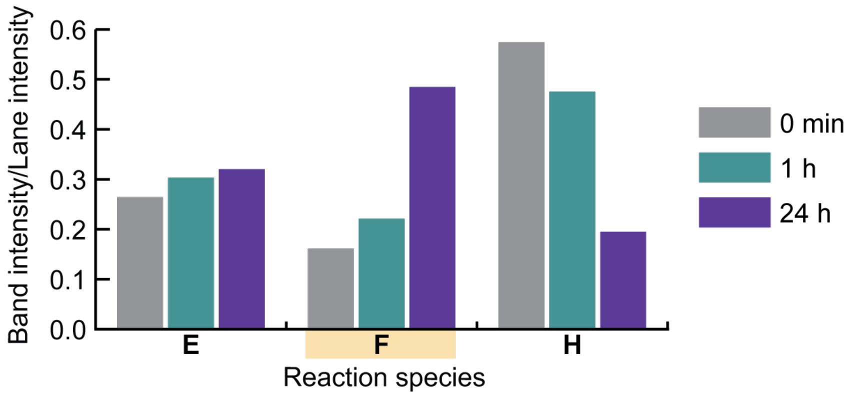


**Figure S5.** Quantitation of gel band intensities at 0 min, 1 h, and 24 h. The formation of GFP-WALP, **F**, product is highlighted in yellow. The depletion of the intermediate species, **H**, is also followed through time.

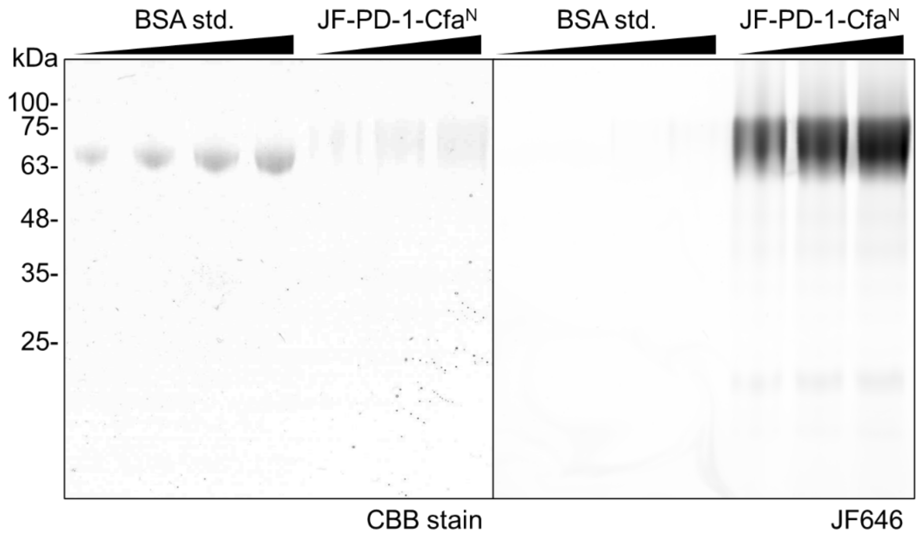


**Figure S6.** SDS-PAGE gel after labeling and purifying JF-PD-1-$Cfa^N$ (51.0 kDa). BSA is used as a control and standard for PD-1 quantification.

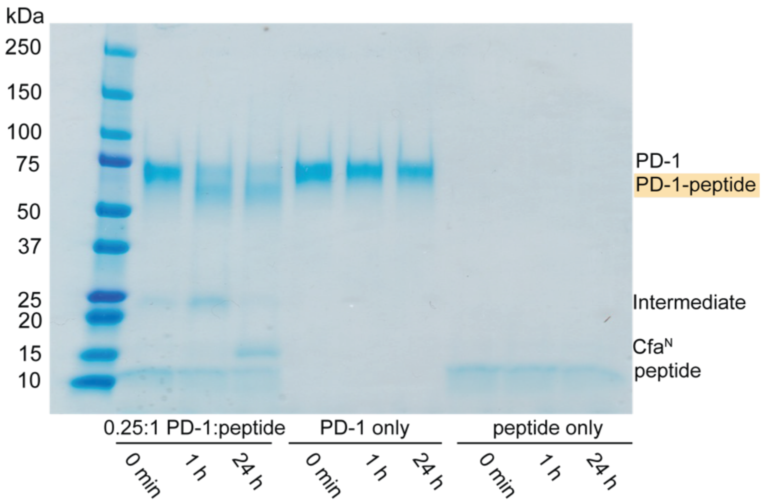


**Figure S7.** SDS-PAGE gel of the reaction between JF-PD-1-Cfa$^{N}$ and Cfa$^{C}$-WALP and controls (JF-PD-1-Cfa$^{N}$ and Cfa$^{C}$-WALP alone). Product is highlighted in yellow.

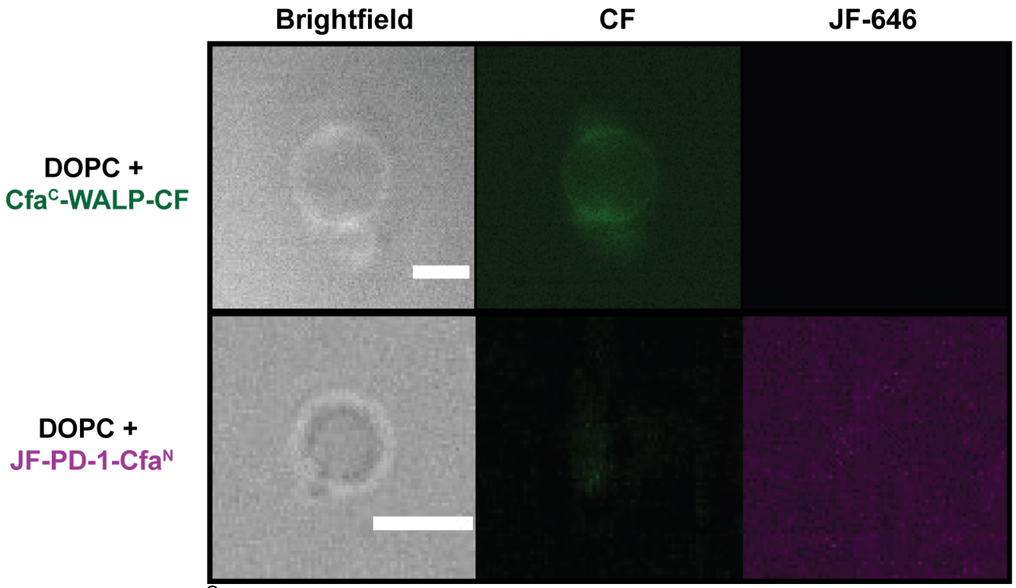


**Figure S8.** Cfa$^{C}$-WALP-CF fluorescence localizes with a brightfield image of peptide reconstituted GUVs and the CF signal does not show up in the JF-646 signal (top row). The bottom row shows the JF-PD-1-Cfa$^{N}$ signal does not localize at the membrane without the presence of Cfa$^{C}$-WALP-CF. Scale bar, 10 μm. CF = carboxyfluorescein. JF-646 = Janelia Fluor 646.

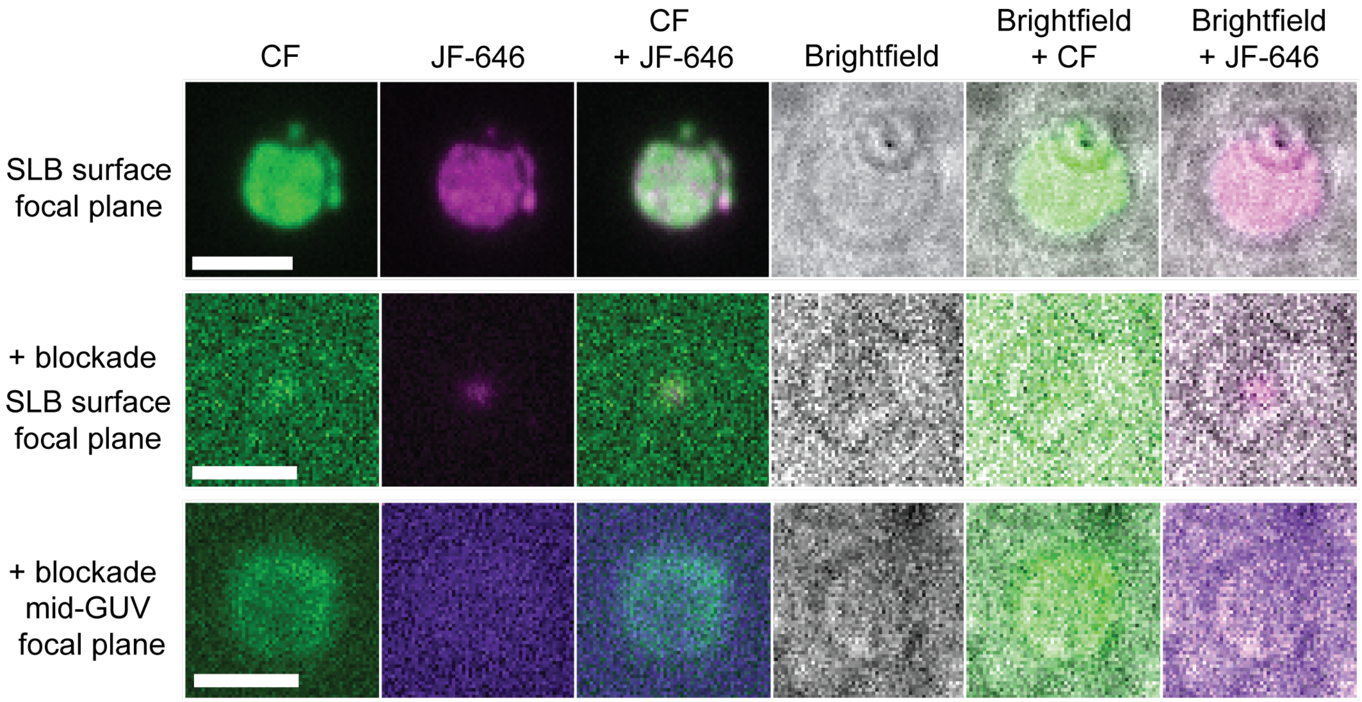


**Figure S9.** Controls and leveled TIRF microscopy images of PD-1/PD-L1 microcluster formation on SLBs +/- blockade. Images are taken at both the SLB surface and mid-GUV focal plane for the + blockade condition to verify the presence of a non-binding GUV above the SLB surface. CF = carboxyfluorescein. JF-646 = Janelia Fluor 646.

### 5. LCMS Spectra

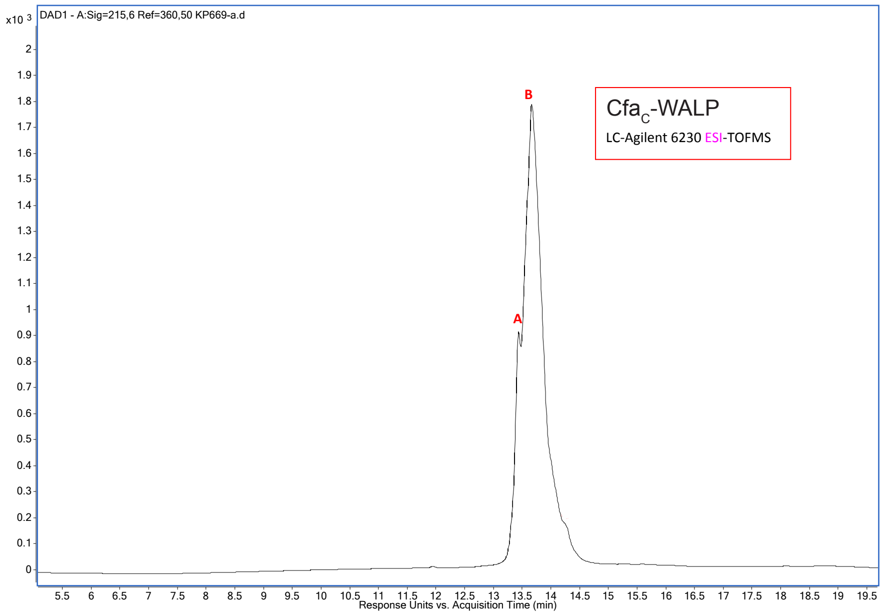


A

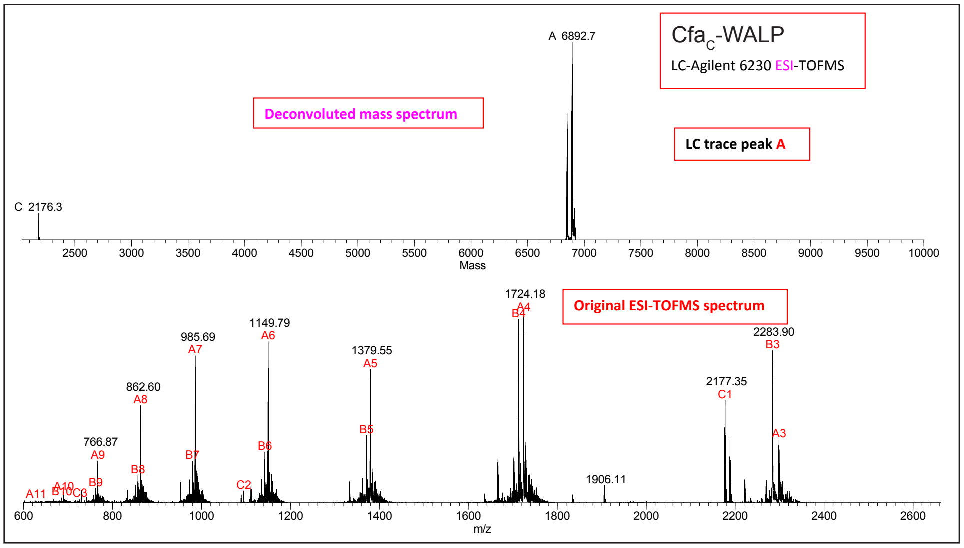


B

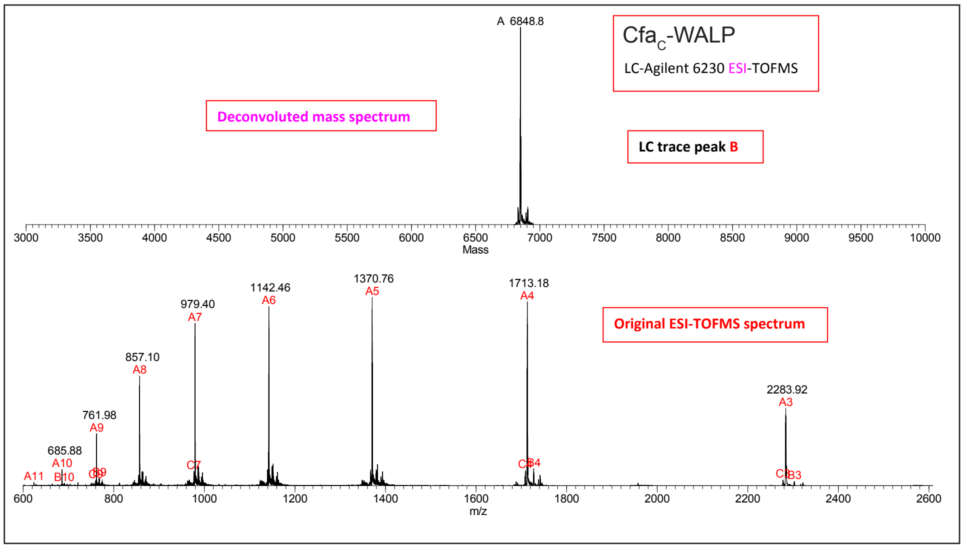


**LC-ESI-MS Spectra 1.** ESI-TOFMS and deconvoluted mass spectra of a LC run of Cfa$^C$-WALP. Top panel is the LC trace of the sample (DAD 215 nm) while subsequent panels are the ESI-TOFMS and deconvoluted mass spectra of the corresponding labeled peaks of the LC trace. Predicted mass: 6849.1 Da.

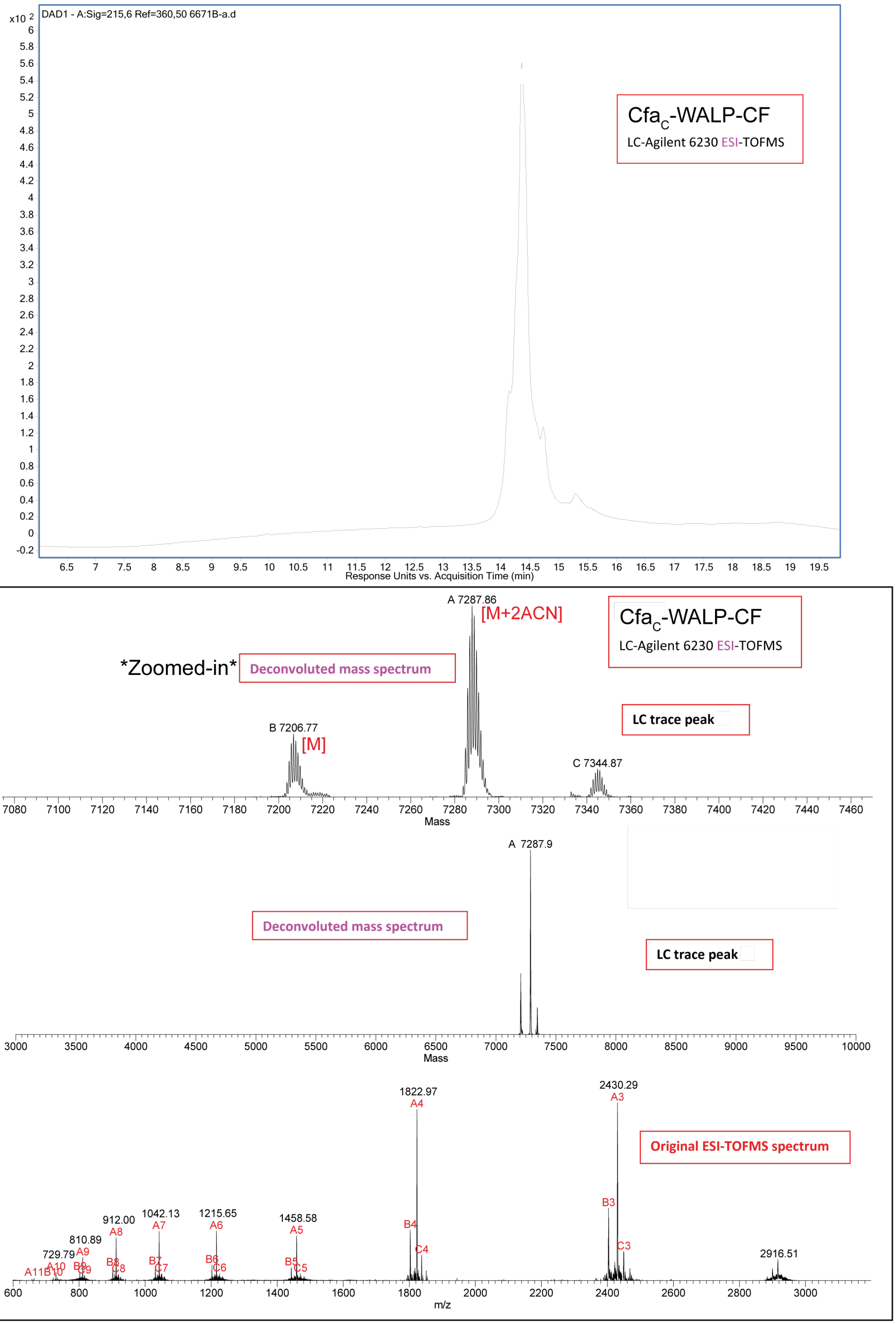


**LC-ESI-MS Spectra 2.** ESI-TOFMS and deconvoluted mass spectra of a LC run of Cfa$^{C}$-WALP-CF. Top panel is the LC trace of the sample (DAD 215 nm) while subsequent panels are the ESI-TOFMS and deconvoluted mass spectra of the corresponding labeled peaks of the LC trace. Predicted mass: 7207 Da.

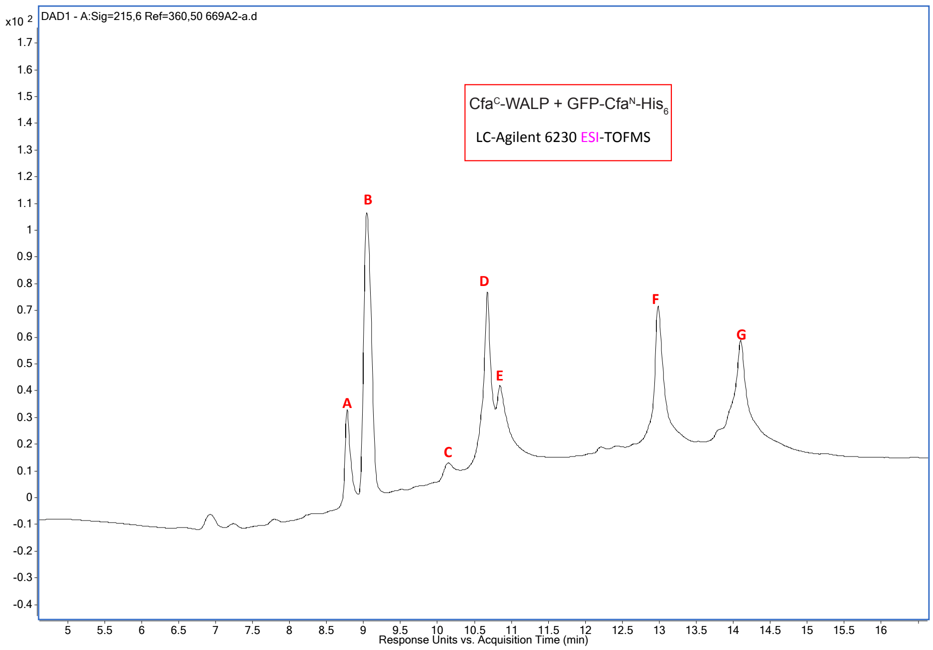
DAD1 - A:Sig=215,6 Ref=360,50 669A2-a.d
x10 2
CfaC-WALP + GFP-CfaN-His6
LC-Agilent 6230 ESI-TOFMS
A
B
C
D
E
F
G
Response Units vs. Acquisition Time (min)

A

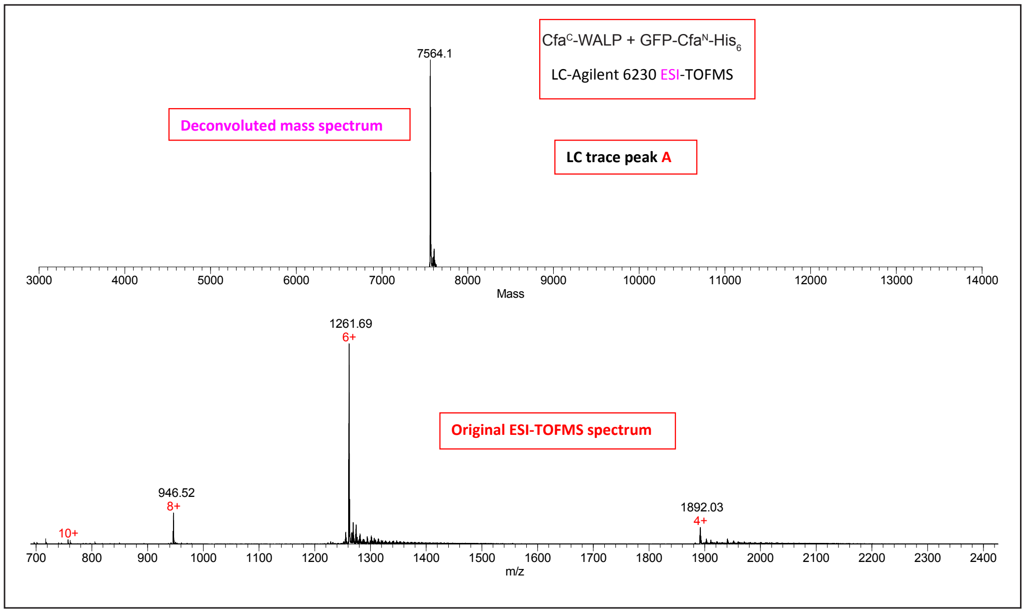
CfaC-WALP + GFP-CfaN-His6
LC-Agilent 6230 ESI-TOFMS
7564.1
Deconvoluted mass spectrum
LC trace peak A
Mass
1261.69
6+
Original ESI-TOFMS spectrum
946.52
8+
10+
1892.03
4+
m/z

B

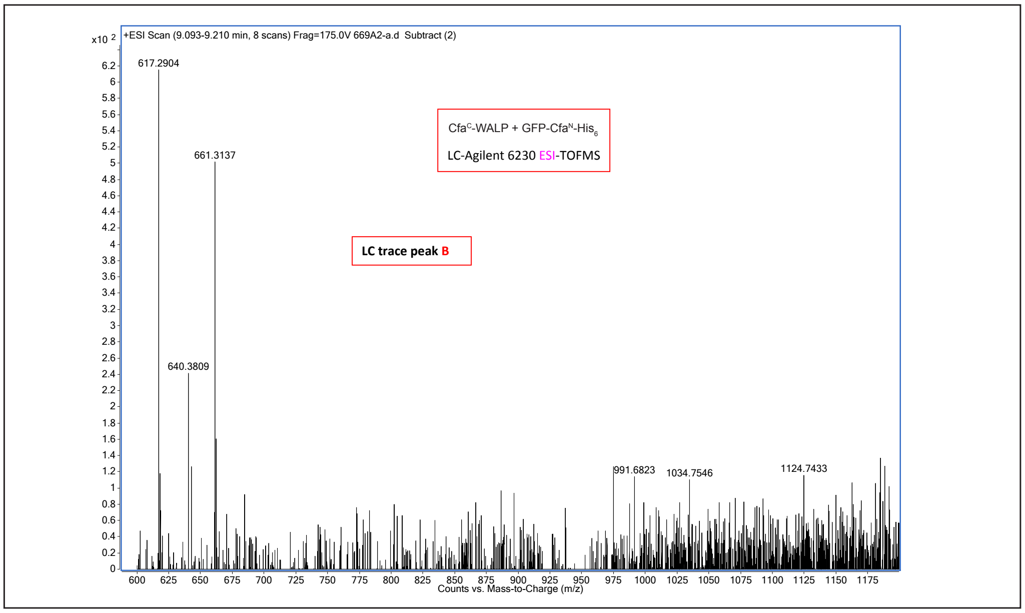
+ESI Scan (9.093-9.210 min, 8 scans) Frag=175.0V 669A2-a.d Subtract (2)
x10 2
617.2904
661.3137
640.3809
CfaC-WALP + GFP-CfaN-His6
LC-Agilent 6230 ESI-TOFMS
LC trace peak B
991.6823
1034.7546
1124.7433
Counts vs. Mass-to-Charge (m/z)

C

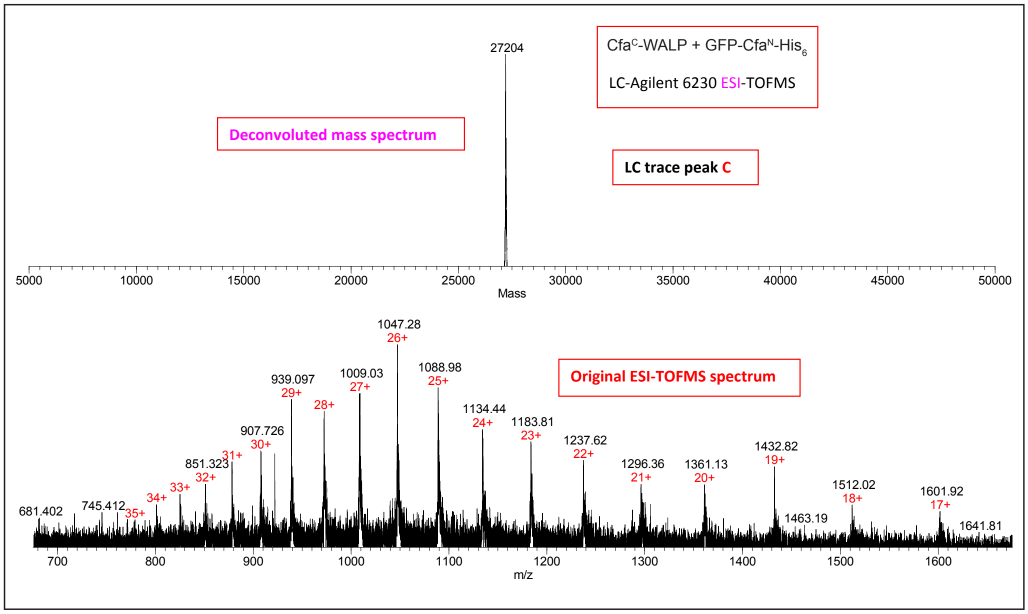
Cfa^C-WALP + GFP-Cfa^N-His6
LC-Agilent 6230 ESI-TOFMS
Deconvoluted mass spectrum
LC trace peak C
27204
5000 10000 15000 20000 25000 30000 35000 40000 45000 50000
Mass
Original ESI-TOFMS spectrum
1047.28 26+
1088.98 25+
1009.03 27+
939.097 29+
28+
907.726 30+
31+
851.323 32+
33+
34+
745.412 35+
681.402
1134.44 24+
1183.81 23+
1237.62 22+
1296.36 21+
1361.13 20+
1432.82 19+
1463.19
1512.02 18+
1601.92 17+
1641.81
700 800 900 1000 1100 1200 1300 1400 1500 1600
m/z

D

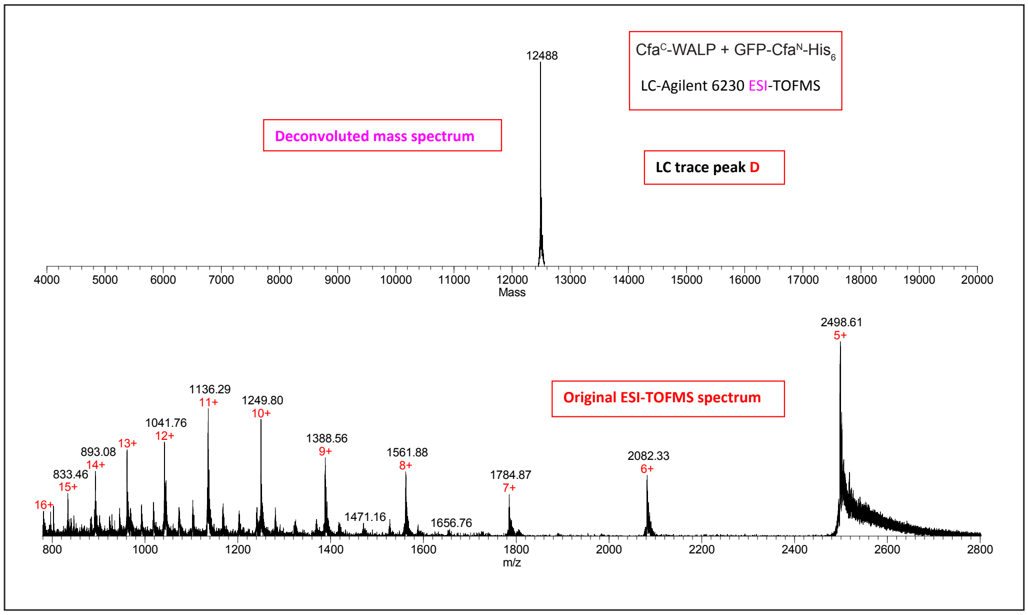
Cfa^C-WALP + GFP-Cfa^N-His6
LC-Agilent 6230 ESI-TOFMS
Deconvoluted mass spectrum
LC trace peak D
12488
4000 5000 6000 7000 8000 9000 10000 11000 12000 13000 14000 15000 16000 17000 18000 19000 20000
Mass
Original ESI-TOFMS spectrum
2498.61 5+
1136.29 11+
1249.80 10+
1041.76 12+
13+
893.08 14+
833.46 15+
16+
1388.56 9+
1471.16
1561.88 8+
1656.76
1784.87 7+
2082.33 6+
800 1000 1200 1400 1600 1800 2000 2200 2400 2600 2800
m/z

E

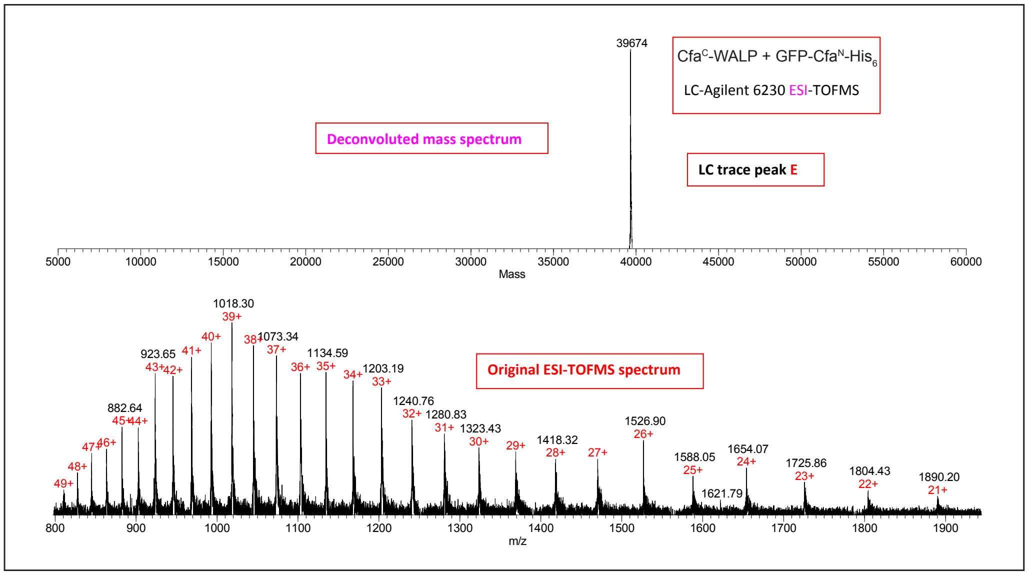
Cfa^C-WALP + GFP-Cfa^N-His6
LC-Agilent 6230 ESI-TOFMS
Deconvoluted mass spectrum
LC trace peak E
39674
5000 10000 15000 20000 25000 30000 35000 40000 45000 50000 55000 60000
Mass
Original ESI-TOFMS spectrum
1018.30 39+
40+
38+ 1073.34 37+
41+
923.65 43+ 42+
1134.59
36+ 35+
34+ 1203.19 33+
1240.76 32+
1280.83 31+
1323.43 30+
29+
1418.32 28+
27+
1526.90 26+
882.64 45+ 44+
47+ 46+
48+
49+
1588.05 25+
1621.79
1654.07 24+
1725.86 23+
1804.43 22+
1890.20 21+
800 900 1000 1100 1200 1300 1400 1500 1600 1700 1800 1900
m/z

F

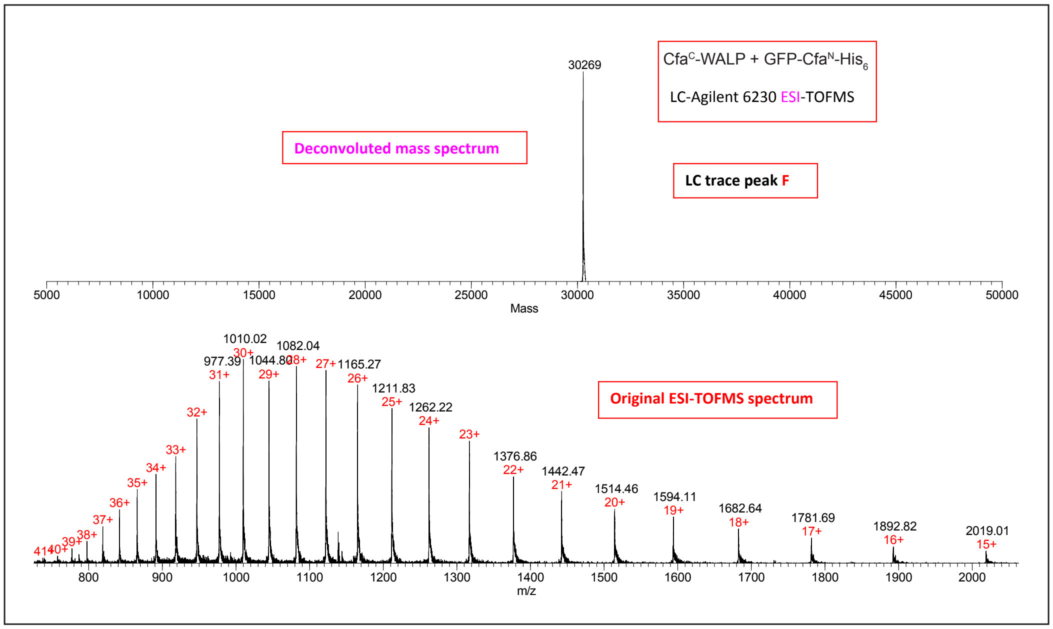


G

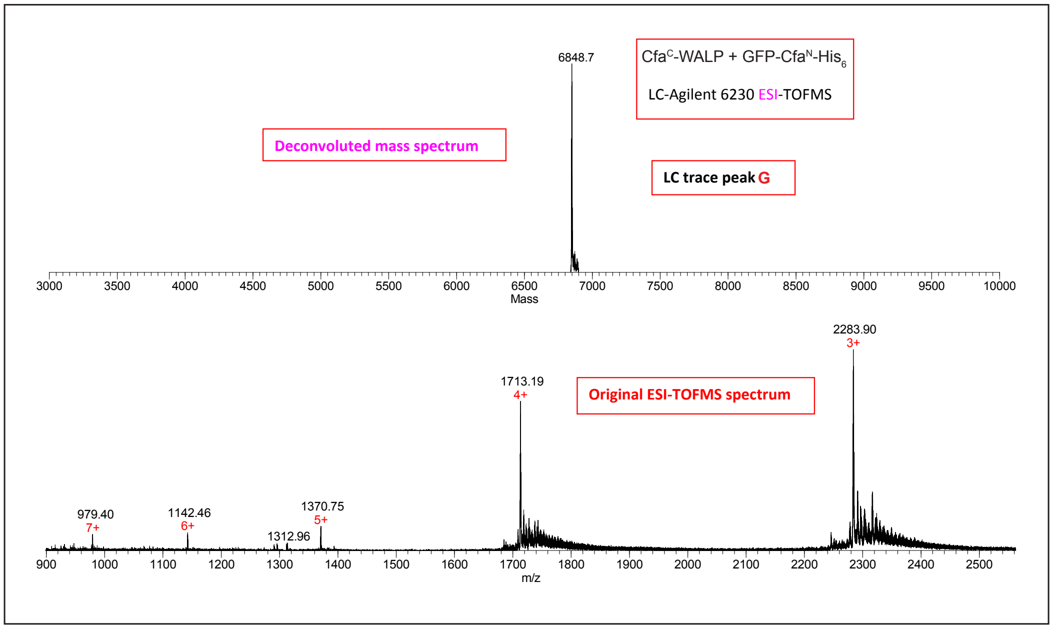


**LC-ESI-MS Spectra 3.** ESI-TOFMS and deconvoluted mass spectra of a LC run of the $Cfa^{C}$-WALP + GFP-$Cfa^{N}$-$His_6$ reaction at 24 h. Top panel is the LC trace of the sample (DAD 215 nm) while subsequent panels are the ESI-TOFMS and deconvoluted mass spectra of the corresponding labeled peaks of the LC trace.
